# Production and functional verification of 8-gene (GGTA1, CMAH, β4GalNT2, hCD46, hCD55, hCD59, hTBM, hCD39)-edited donor pigs for xenotransplantation

**DOI:** 10.1101/2024.05.19.594889

**Authors:** Jiaoxiang Wang, Kaixiang Xu, Tao Liu, Heng Zhao, Muhammad Ameen Jamal, Gen Chen, Xiaoying Huo, Chang Yang, Deling Jiao, Taiyun Wei, Hanfei Huang, Hongfang Zhao, Jianxiong Guo, Fengchong Wang, Xiong Zhang, Kai Liu, Siming Qu, Gang Wang, Hong-Ye Zhao, Zhong Zeng, Hong-Jiang Wei

**Author notes:** Correspondence: Hong-Ye Zhao; Zhong Zeng; Hong-Jiang Wei. Authors contributed equally to this work (Jiaoxiang Wang, Kaixiang Xu, Tao Liu, Heng Zhao, Muhammad Ameen Jamal, Gen Chen).

## Abstract

Gene-edited pig-to-human xenotransplantation continues to make breakthroughs and is expected to enter clinic to solve the global shortage of donor organs. However, which gene combination is suitable for which organ transplantation remains unclear. In this study, we utilized CRISPR/Cas9 gene editing technology, PiggyBac transposon system and somatic cell cloning to construct GTKO/CMAHKO/β4GalNT2KO/hCD46/hCD55/hCD59/hCD39/hTBM 8 gene-edited cloned (GEC) donor pigs, and performed pig to non-human primate (NHP) transplantation to evaluate the effectiveness of these GEC pigs. The multiple vectors were co-transfected into fetal fibroblasts of Diannan miniature pig with O blood type, and 25 colonies were screened out, and one of them carried GGTA1, CMAH and β4GalNT2 biallelic knockout and integration of hCD46, hCD55, hCD59, hTBM and hCD39 genes, which was used as a donor cell for cloning, and a 33-day-old viable fetus was obtained. The fetus was identified and confirmed for normal karyotype and the absence of three xenogeneic antigens α-Gal, Neu5Gc and Sda, and expression of hCD46, hCD55, hCD59, hTBM and hCD39 genes, then the recloning was carried out and 28 cloned piglets were obtained by natural delivery. Molecular identification at DNA, mRNA and protein levels showed that 8 gene editing (GE) was successful in these GEC piglets. Moreover, antigen-antibody binding assay and complement-dependent cytotoxicity assay demonstrated that 8GE effectively reduced the immune incompatibility and kidney xenograft survived up to 15 and 17 days into two NHPs, respectively. During this period, the recipient serum antibodies IgA and IgM, complements C3 and C4, coagulation indicators PT, APTT, TT and FIB, as well as most electrolytes and liver function indicators remained relatively stable. The 24-hour urine output and serum creatinine remained normal at a period of post-transplantation. These results indicated that the 8GEC pigs effectively alleviated immune rejection and exerted life-supporting kidney function in the recipient.

## INTRODUCTION

Xenotransplantation is an effective way to solve the global donor shortage. With the rapid development of gene editing technology and new immunosuppressant, the United States of America (USA) has taken a leading step in conducting subclinical research on gene-edited pig-human xenotransplantation. In 2022, the world’s first GE pig heart was transplanted into a human body at the University of Maryland Medical Center in the United States, which survived for 60 days (Griffith and Goerlich, 2022). In 2023, a kidney graft from a GE pig to a brain-dead patient was transplanted at New York University Langone Medical Center in the USA. This kidney functioned normally for 61 days, thus establishing a new record for functional GE pig kidneys in humans. In recent years, pig-to-human xenotransplantation has made new breakthroughs and is soon expected to enter clinical phase.

The major barriers in pig-to-human organ transplantation are immunological incompatibility, physiological dysfunction and biological safety. The glycogen α-Gal, Neu5Gc and Sda synthesized by three proteins GGTA1, CMAH and β4GalNT2, respectively are known xenoantigens. Once the pig xenograft is transplanted into the recipient, the corresponding natural antibodies binds to these carbohydrate antigens on the surface of the cells, and induces a hyperacute rejection (HAR). Therefore, inactivation of this xenoantigens is necessary. Besides this, there may be/are multiple unknown xenoantigens in pigs. After antigen presentation and B cell activation, corresponding antibodies may still be produced, thereby activating the complement system to attack pig cells. In addition, the development of thrombotic microangiopathy becomes increasingly evident with long-term graft survival (Cowan and Robson, 2015). Therefore, adding complement regulatory proteins (CD46, CD55, CD59) and coagulation regulatory factors thrombomodulin (CD39, TBM) can effectively improve these immune responses (Cozzi and White, 1995; McCurry et al., 1995; Pierson et al., 2009).

In this regard multiple gene-edited (GE) donor pigs with different gene combinations were constructed, and transplanted to non-human primates, and demonstrated their effectiveness in long-term survival of xenograft. In pig-to-nonhuman primate (NHP) xenotransplantation, transplantation of pig heart with 3-GE (GTKO/hCD46/hTBM) into baboon functionally survived up to 195 days (Langin et al., 2018). The kidneys from 2-GE (GTKO/hCD55) pig functionally survived up to 499 days in rhesus monkeys (Kim et al., 2019). Recently, the transplantation of 10 gene-edited pig kidneys into cynomolgus monkeys extended the survival up to 758 days (Anand et al., 2023). Furthermore, two cases of recent heart transplant and multiple cases of kidney transplantation from GE pig to a human patient with organ failure or brain-dead humans exhibited the feasibility of donor pig to save human lives (Griffith et al., 2022; Moazami et al., 2023). However, at present, it is extremely difficult to edit all these genes related to xenorejection and obtain viable pigs, thus only selective gene-editing could be performed, and however, which genetic combination is most suitable for which organ remains undetermined. Thus, there is a need to utilize the different genetic combinations and gene editing strategies for development of the GE donor pigs of for different xenograft and to ensure the long-term survival of xenograft in preclinical and clinical xenotransplantation.

This study used Cas9, PiggyBac transposon and somatic cell nuclear transfer (SCNT) technology to construct 8-GE (GTKO/CMAHKO/β4GalNT2KO/hCD46/hCD55/hCD59/hTBM/hCD39) *Diannan* miniature pigs and performed kidney transplantation from pig to rhesus monkey to evaluate the effectiveness of these GE donor pigs.

## RESULTS

### Generation of the 8-GEC pigs

To produce the GTKO/CMAHKO/β4GalNT2KO/hCD46/hCD55/hCD59/hTBM/hCD39 8-GEC donor pigs, the CRISPR/Cas9 system, the PiggyBac transposons system and somatic cell cloning were applied and the constructed plasmids were co-transfected into fetal fibroblasts of *Diannan* miniature pigs. After drug selection and genotyping, 8-GE cell colonies were obtained and used as donor for somatic cell cloning to obtain 8-GEC pigs (Figure 1A). We designed two sgRNAs targeting the 3^rd^ exon of GGTA1 gene, two sgRNAs targeting the 4^th^ exon of CMAH gene, and 3 sgRNAs targeting the 2^nd^ exon of β4GalNT2 genes to simultaneously KO these three genes in the *Diannan* miniature pigs (Figure 1B). The overexpression of hCD46, hCD55, hCD59, hTBM and hCD39 genes was achieved using the human ICAM-2 promoter (hTBM and hCD39), and the human EF-1α promoter (hCD46, hCD55 and hCD59) (Figure 1C). After electroporation and drug selection, a total of 25 cell colonies were obtained. At first, these colonies were identified by PCR and 64% (16/25) carried humanized genes (Figure 1D). Then, 8 colonies were randomly selected for Sanger sequencing and results showed that 7 colonies (C2#, C3#, C5#, C8#, C11#, C12# and C21#) for GGTA1, 5 colonies (C2#, C5#, C8#, C11# and C12#) for CMAH, and 1 colony (C12#) for β4GalNT2 were biallelic KO. Therefore, among the 25 colonies, only the C12# colony was biallelic KO of 3 endogenous genes along with integration of 5 humanized genes (Figure 1E, Table 1), thus it was used as a donor for SCNT and reconstructed embryos were transferred into 4 surrogate sows. One of them became pregnant and a 33-day-old viable fetus (C12F01) was obtained (Figure 2A, Table 2), which also carried 5 humanized genes along with biallelic KO 3 genes as verified by genotyping (Figure 2B, C) and exhibited normal karyotype (Figure 2D). Moreover, immunofluorescence analysis also showed that three xenoantigens αGal, Neu5Gc and Sda were deficient, and hCD46, hCD55, hCD59, hTBM and hCD39 genes were expressed in the fetus, (Figure 2E). Therefore, we used this fetal fibroblasts as donor cells for re-cloning and transferred them into 17 surrogate sows. Six of them became pregnant, giving birth to a total of 28 piglets, of which 24 survived with an average of 1.4 live pigs per sow (Table 3). Out of the surviving piglets, 17 grew up healthy and reached to sexual maturity (Figure 4A, Table 4). So far, some have been cross-bred to produce viable F1-generation individuals.

**Figure 1.**
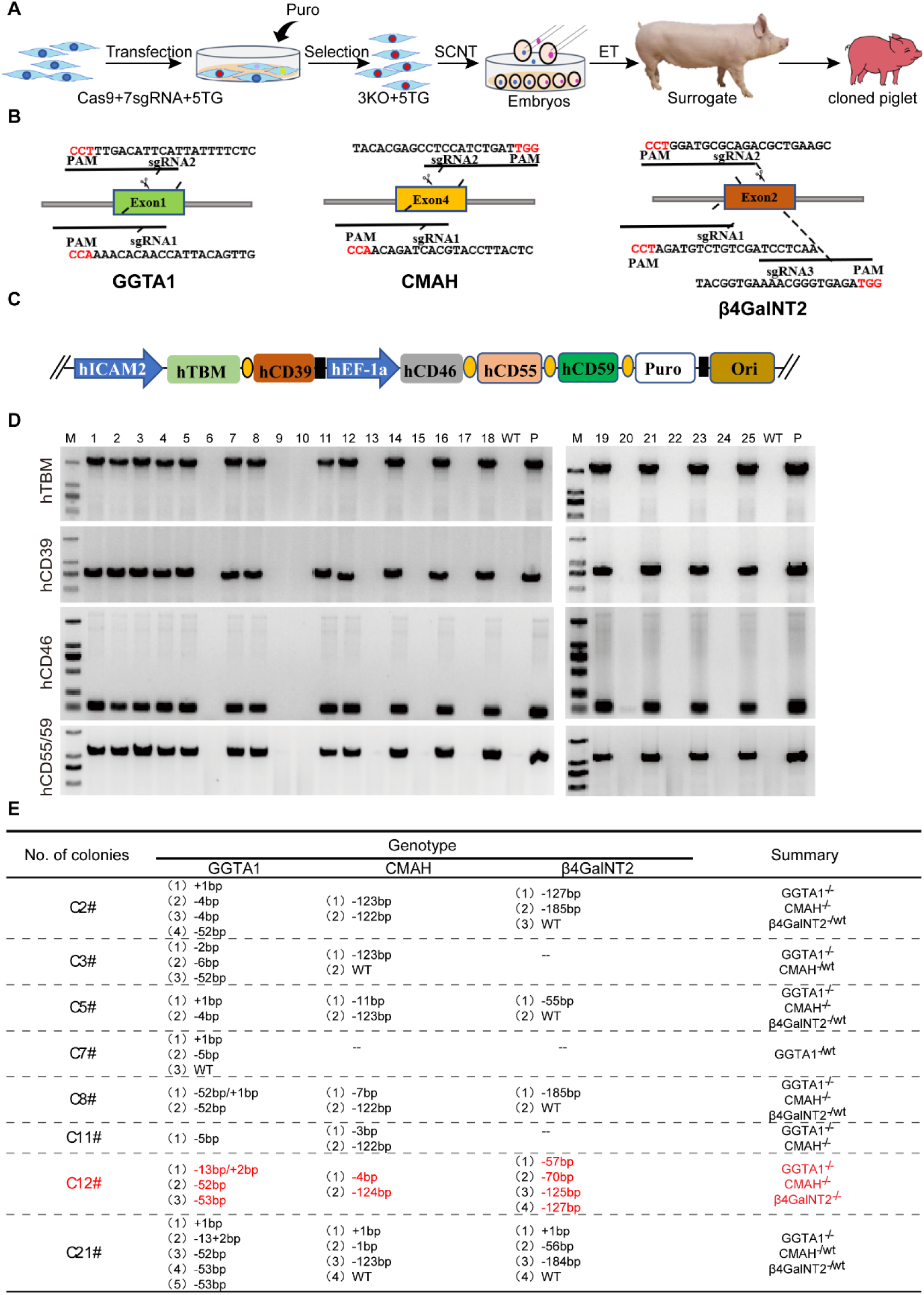
Construction of gene editing vectors for 3 gene knockout, and 5 genes overexpression as well as screening out of cell colonies. **A.** Schematic diagram for the generation of 8-GE cloned piglets. **B.** Knockout of porcine GGTA1, CMAH, β4GalNT2 gene by CRISPR/Cas9 targeting Exon 3, 4 and 2, respectively. C. The expression of cDNA of human CD46, hCD55 and hCD59 genes was driven by human EF-1α promoter, and the expression of cDNAs of hTBM and hCD39 gene were driven by human ICAM2 promoter. **D.** hTBM, hCD39, hCD46, hCD55 and hCD59 genes were successfully intergrated into pig genome in cell colonies by PCR identification. A total of 16 cell colonies were positive for transgene integration. **E.** The sequence of the targeting region of GGTA1, CMAH, β4GalNT2 genes in cell colonies by Sanger sequencing after transfection and drug selection. Only the C12# colony was biallelic KO of 3 endogenous genes along with integration of 5 humanized genes.

**Figure 2.**
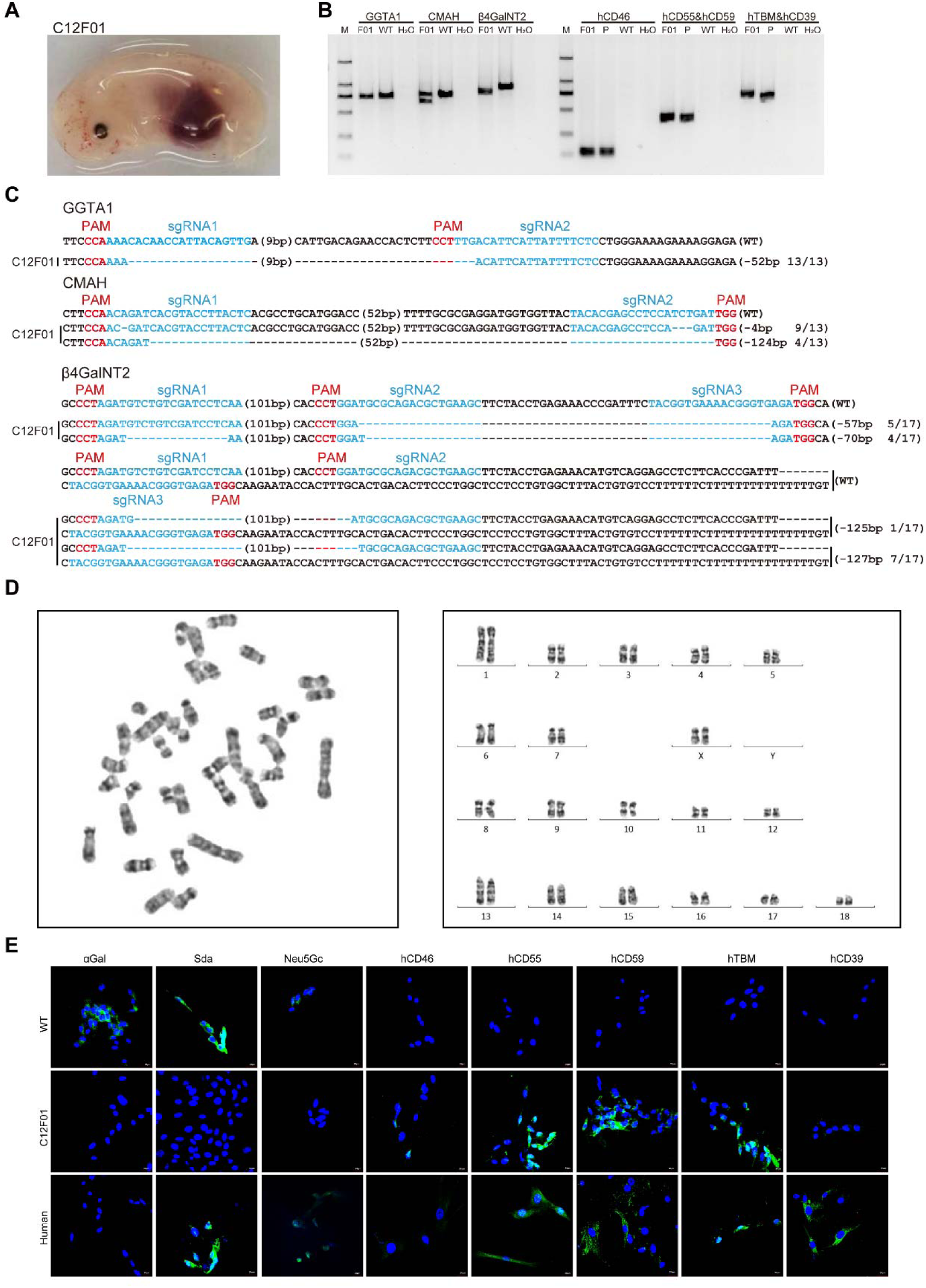
Generation and identification of gene editing cloned fetuses. **A.** Photo of cloned fetus (C12F01) obtained after 33-day pregnancy. **B.** The biallelic KO of 3 genes and integration of 5 genes genes into fetal genome was confirmed by PCR. **C.** The sequence of the targeting region of GGTA1, CMAH, and β4GalNT2 genes in fetus by Sanger sequencing. **D**. Karyotying analysis of cloned fetus. **E**. The protein expression of αGal, Neu5Gc, Sda, hCD46, hCD55, hCD59, hTBM and hCD39 genes in cloned fetus was confirmed by immunofluroscence.

### Phenotypic identification of 8-GEC pigs

We determined the genotype of targeting region at GGTA1, CMAH, and β4GalNT2 genes using Sanger sequencing and results showed that the cloned pigs were biallelic knockouts for all 3 genes at genomic levels. Among them, the GGTA1 gene has a 52bp, the CMAH gene has 4bp, and 124bp deletion mutations, and the β4GalNT2 gene has 57bp, 70bp, 125bp and 127bp deletion mutation, which were consistent with the C12F01 fetal genotype (Table 5). Moreover, the copy number of 5-transgene (hCD46, hCD55, hCD59, hTBM and hCD39) in heart, liver, lung, kidney, and spleen tissues of cloned piglets were maintained at 2∼3 copies, while those of human cells and pre-transfected control cells was 2 and 0 copies, respectively (Figure 3). Further, results of mRNA expression levels showed that all transgenes were expressed in the heart, kidney, liver, and lung tissues of cloned piglets. The mRNA expression levels of hCD46 and hCD39 were higher, while of hCD55, hCD59 and hTBM were lower in the kidney of 8-GEC pigs. It is worth noting that all genes exhibited higher expression in lung tissues, while lower expressions in heart tissue (Figure 4B-F). The immunofluorescence analysis showed that αGal, Neu5Gc and Sda were deficient, while hCD46, hCD55, hCD59, hTBM and hCD39 were expressed in the kidney tissue of cloned piglets. Among them, the protein expression of hCD55, hCD59, hTBM and hCD39 genes in P09 cloned piglets was highly significant (Figure 4G). In addition, we found that the protein expression levels of the same gene in different tissues of the same cloned individual (P03) were not consistent (Figure S1). Nevertheless, when rhesus monkey serum was used to perform antigen-antibody binding, and complement-dependent cytotoxicity assays, the binding ability of monkey IgG and IgM to 8-GEC pigs’ PBMCs was significantly reduced (Figure 4H, I), and complement attack of porcine PBMCs was significantly weakened (Figure 4J). The above results showed that we successfully generated the 8 (GTKO/CMAHKO/β4GalNT2KO/hCD46/hCD55/hCD59/hTBM/hCD39) GEC pigs by using gene editing and somatic cell cloning technology.

**Figure 3.**
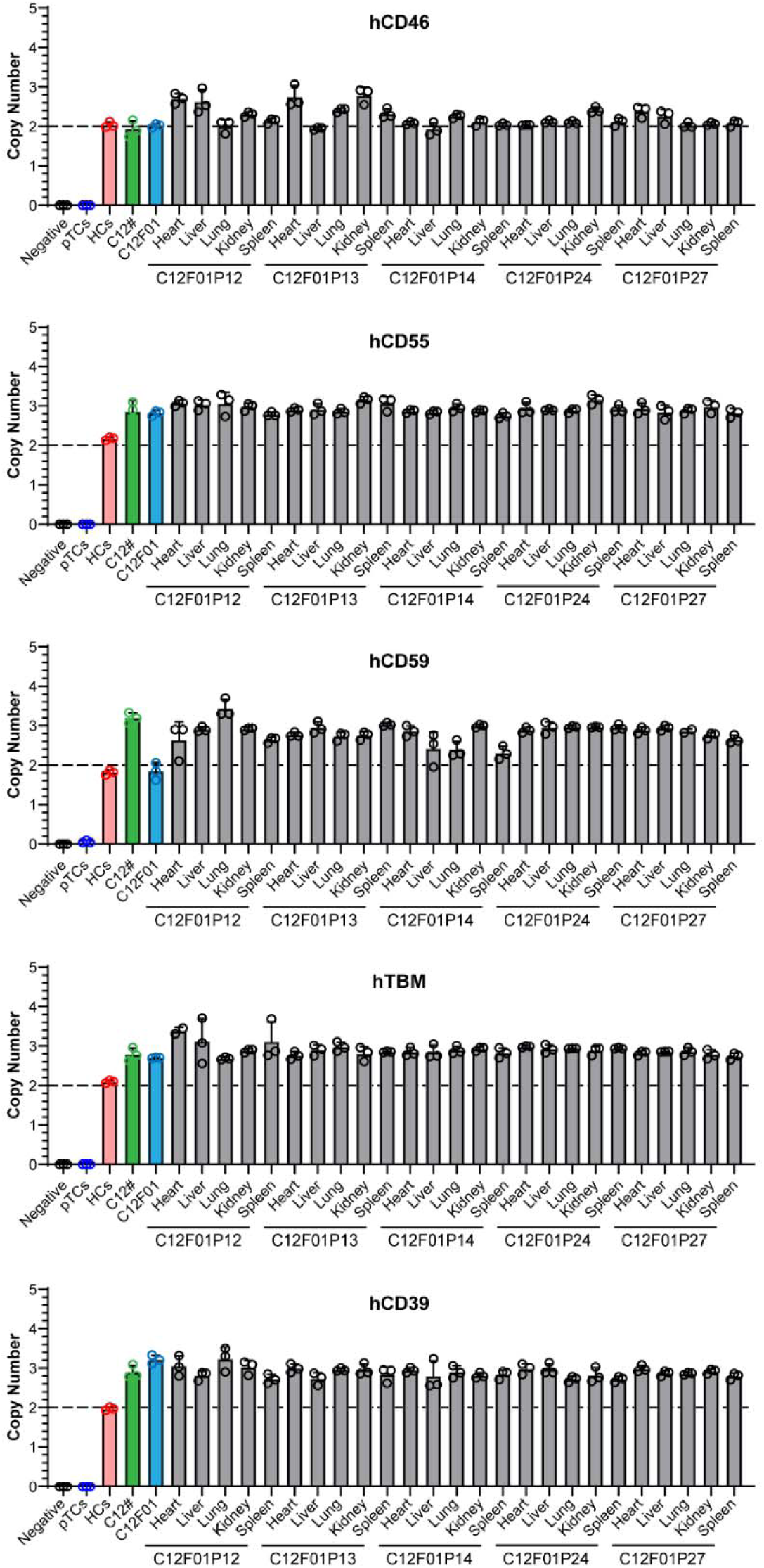
Copy numbers of transgenes hCD46, hCD55, hCD59, hTBM and hCD39 in donor cell lines, cloned fetus, and piglets. The copy numbers of all transgenes were detected by ddPCR. Negative, water; pTCs, porcine pre-transfected cells; WT, wild-type pig kidney; HCs, human cells.

**Figure 4.**
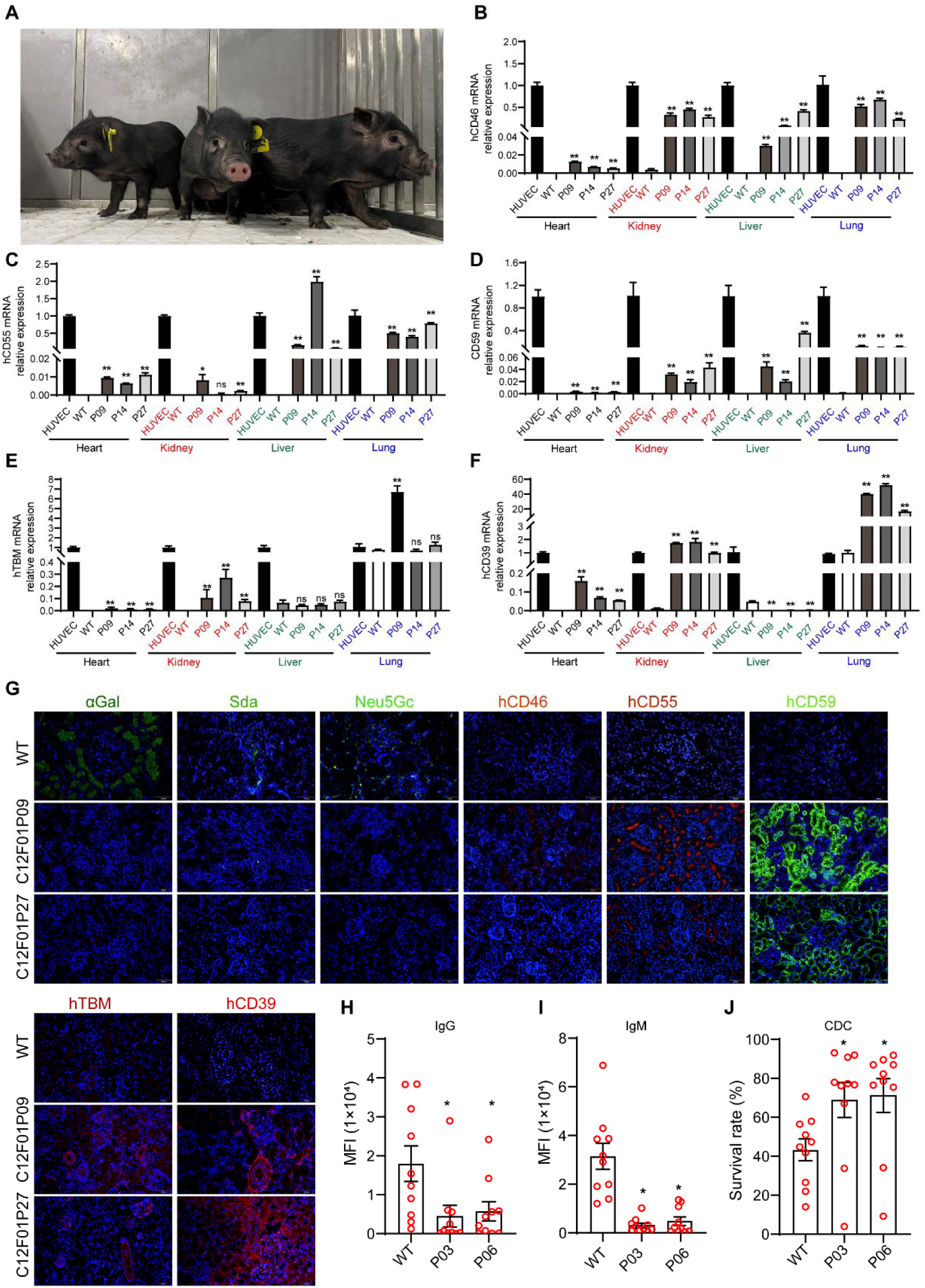
Phenotype of 8-GE cloned pigs and crossmatch with rhesus monkeys. A. Photo of GEC piglets. **B-F**. The mRNA expression levels of hCD46, hCD55, hCD59, hTBM, and hCD39 genes in heart, kidney, liver, and lung tissues of GEC pigs (n=3; P09, P14, P27). WT, wild type; HUVEC, Human umbilical vein endothelial cell. **G**. The expression of α αGal, Neu5Gc, Sda, hTBM, hCD39, hCD46, hCD55 and hCD59 in GEC pig kidney was confirmed by immunofluroscence. **H**. The levels of monkey IgG binding to 8-GE porcine peripheral blood mononuclear cells (PBMCs). **I**. The levels of monkey IgM binding to 8-GE porcine PBMCs. **J**. The survival rate of 8-GEC porcine PBMCs after after incubation with each monkey serum.

### Kidney xenotransplantation from 8-GEC pigs to non-human primates

In order to evaluate the effectiveness of 8-GEC pigs as donor pigs for xenotransplantation, we carried out two cases of pig-to-monkey kidney transplantation. First, we collected the serum from 7 monkeys and conducted cross-matching experiment with the PBMCs of two 8-GC pig (P03 and P06) and selected 2 monkeys (R24#, R25#) as recipient for kidney xenotransplantation (Figure S2). Two days before surgery, immunosuppression was performed with anti-CD20 mAb and ATG respectively. On the day of the operation, the P03 pig suddenly died during the kidney removal process so we first performed a kidney transplant from P06 pig to the R25# monkey. After removing one kidney of the monkey, the pig kidney was anastomosed to the monkey’s blood vessel *in-situ* and the blood flow was opened. The color of the pig kidney immediately changed from white to pink, and no HAR was found (Figure S3B). Considering that the pig kidney may be difficult to maintain function due to its small size, thus we retained a monkey kidney and completed the transplantation.

Next, we re-selected 8-GEC pig (P23) with a suitable weight and continued to complete the kidney transplantation from pig to monkey R24#. Three days before surgery, immunosuppression was performed with anti-CD20 mAb, ATG and CVF respectively. On the day of surgery, we first removed one monkey kidney, then transplanted the pig kidney to the recipient monkey, and finally removed the other monkey kidney. The transplantation operation was successful. The pig kidney produced urine within minutes after the transplantation, and no hyperacute rejection was found. To facilitate postoperative drug administration, we performed a gastrostomy in the recipient monkeys. During the postoperative care process, regular administration was carried out according to the established immunosuppressive regimen (Figure S3A), and corresponding treatment or nutritional supplements were given accordingly. We monitored the concentrations of drug administered through gastrostomy and found that tacrolimus, MMF, and sirolimus immunosuppressants were not detected in the blood 3 days after surgery. Later, we detected drug concentrations in serum only after replacing oral immunosuppressants with injectable dosage (Figure S3C). The ultrasonography of the recipient monkey on day 6 after surgery showed that the pig kidneys still maintained the full blood flow without intravascular thrombosis (Figure S3D). Unfortunately, two recipients M1 and M2 died on day 17 and 15 post-transplantation.

We summarized the entire pig-to-monkey kidney transplantation process. In terms of immune rejection, the serum antibodies IgA, IgG and IgM, and complements C3 and C4 of the recipient monkeys were maintained at relatively stable levels (Figure 5A and B).

**Figure 5.**
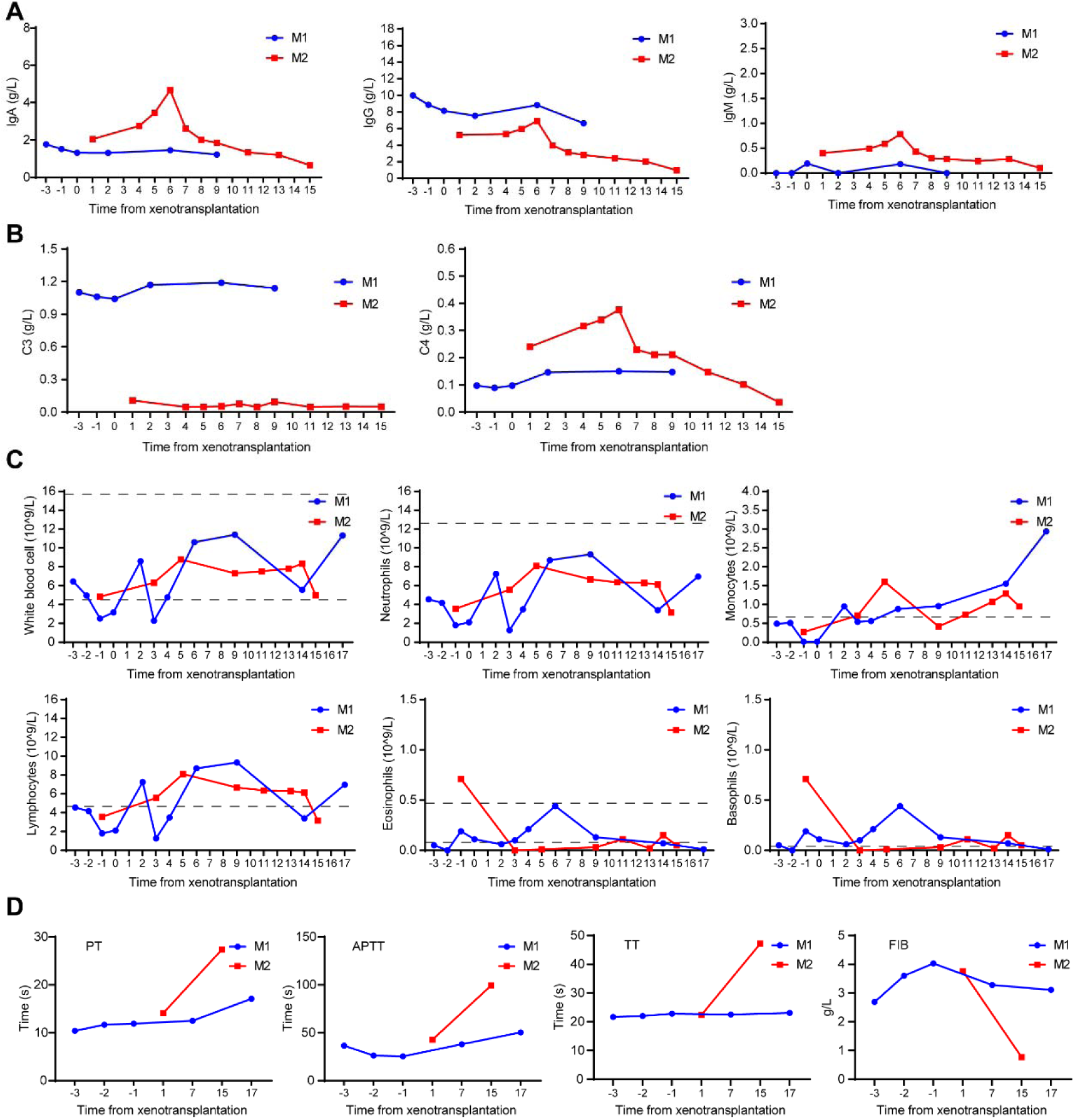
Humoral, cellular immunity and coagulation of recipient monkey. **A.** The levels of IgA, IgG, and IgM in serum of recipient monkey. **B**. The levels of complement C3 and C4 in serum of recipient monkey. **C.** The coagulation indexes: **A**. Thrombin time (TT), prothrombin time (PT), activated partial thromboplastin time (APTT) and fibrinogen (FIB) in recipient monkey. **D.** The numbers of white blood cells(WBC), neutrophils, basophils, eosinophils, lymphocytes and monocytes in the whole blood of recipient monkey.

The number of leukocytes, neutrophils, lymphocytes, eosinophils and basophils was within a controllable range, while the number of monocytes gradually increased during the process (Figure 5C). In terms of coagulation, PT, APTT, TT and FIB remained relatively stable in the recipient M1, but increased significantly the recipient M2 (Figure 5D).

In terms of renal pathology, HE staining revealed a large amount of red blood cell leakage in the renal graft (Figure 6A). Immunofluorescence staining of 8-GEC pig kidney grafts revealed a small amount of IgG and IgM antibodies as well as complement deposition of C4d, C3c, and C5b-C9 in the graft (Figure 6B). In addition, immunohistochemical staining revealed that the graft was infiltrated with CD57^+^ NK cells and CD68^+^ macrophages (Figure 6C). The above results showed that 8-GE pig kidneys can effectively overcome the HAR, but gradually develop acute humoral rejection and cellular immune rejection.

**Figure 6.**
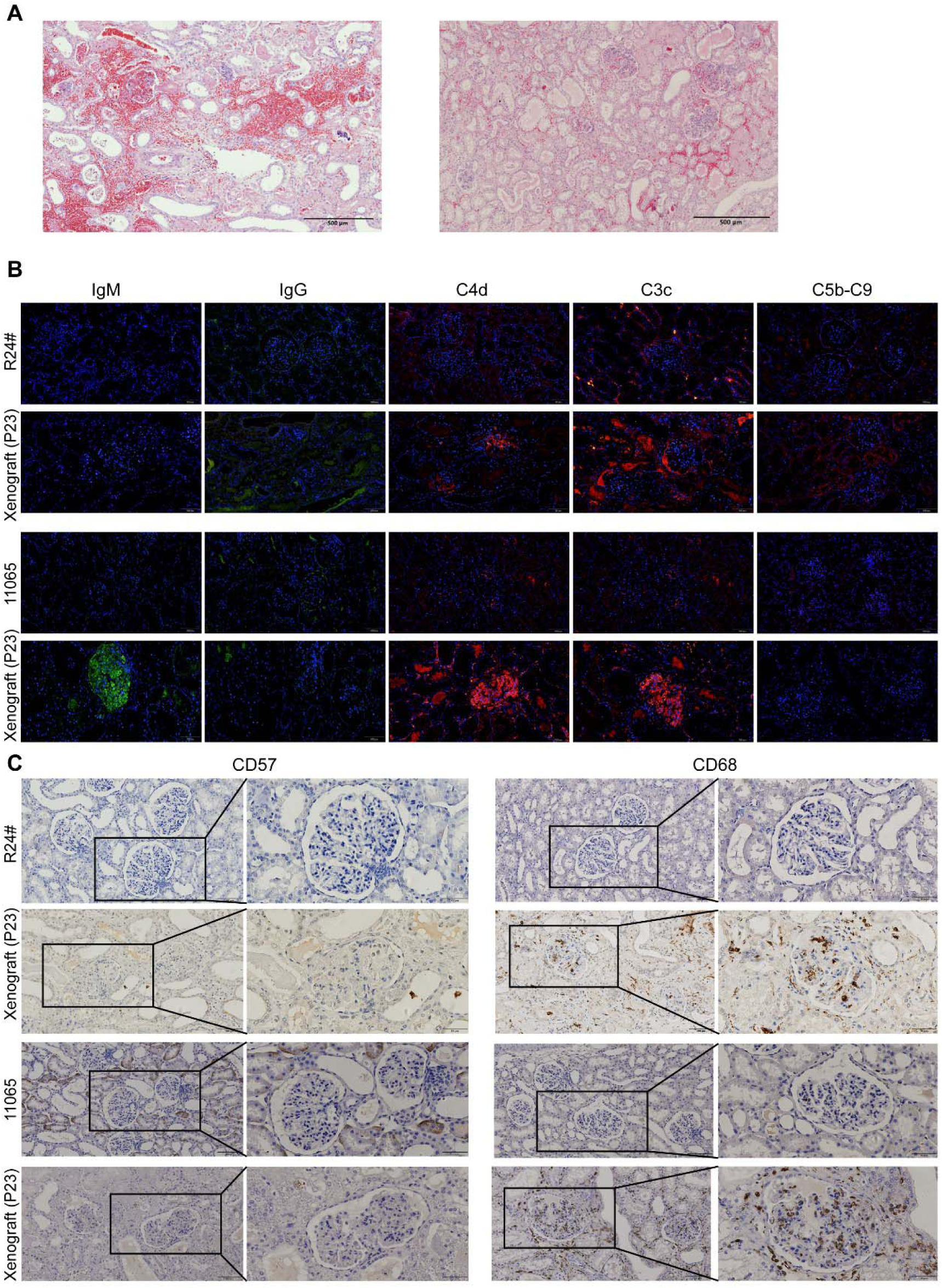
Pathological section of renal tissue. **A.** HE staining of kidney xenograft. **B.** Immunoglobulins and complement deposition in 8-GE porcine kidney xenograft confirmed by immunofluorescence. **C.** NK cells and macrophage infiltration in porcine kidney xenograft confirmed by immunohistochemistry, scale bar=100 μm.

In terms of renal function, the 24-hour urine output of recipient monkeys remained good in the early stages of post-transplantation but gradually decreases in the later stages of post-transplantation (Figure 7A). The serum creatinine levels remained stable in the recipient M1 but fluctuated in the recipient M2 (Figure 7B). The serum urea level also remained stable in the recipient M1 but was obviously elevated in the recipient M2(Figure 7C). While, the other indicators such as serum uric acid and cystatin C were maintained at relatively stable levels (Figure 7D and E), indicating that the kidney function of the 8-GE pigs was basically normal. The kidney also promoted the production of RBCs and therefore, we monitored the changes in the number of RBCs. It was found that RBCs showed a gradual decreasing trend (Figure 7F). In addition, both of recipients developed severe thrombocytopenia (Figure 7G).

**Figure 7.**
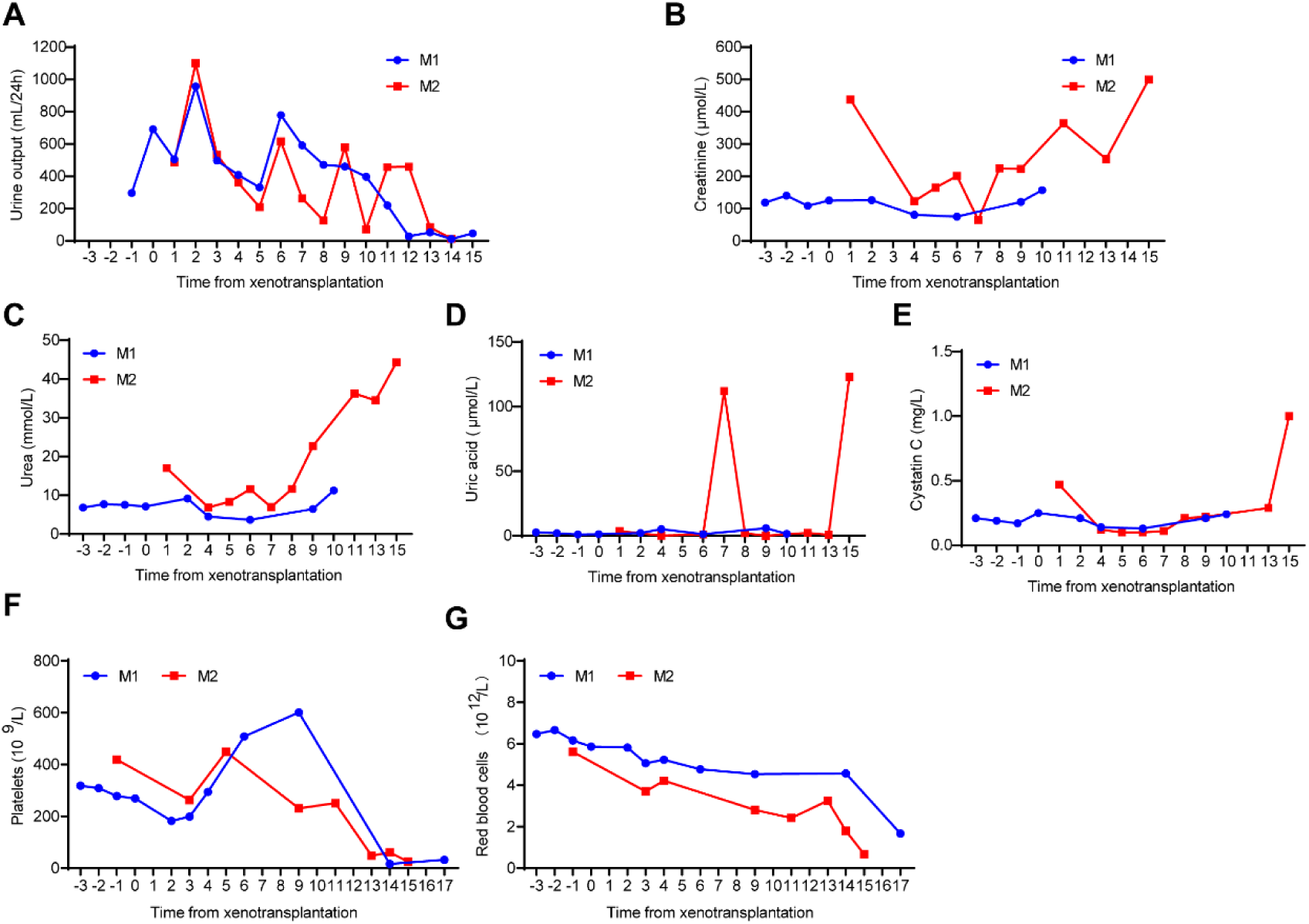
The renal function and erythrocytes indexes of recipient monkey. **A.** 24-hour urine output. **B-E.** The levels of creatinine (B), urea (C), uric acid (D) and cystatin C (D) in the serum of recipient monkey. **F-G**. The numbers of platelets (F) and red blood cells (G).

In terms of liver and other physiological functions, various liver function indicators, including TP, ALB, GLB, TBIL, TBA, AST, ALT, ADA and ALP remained relatively stable (Figure 7C-E). Electrolytes Na, Cl, HCO_3_, Mg, P, Ca and K also remained stable (Figure 8).

**Figure 8.**
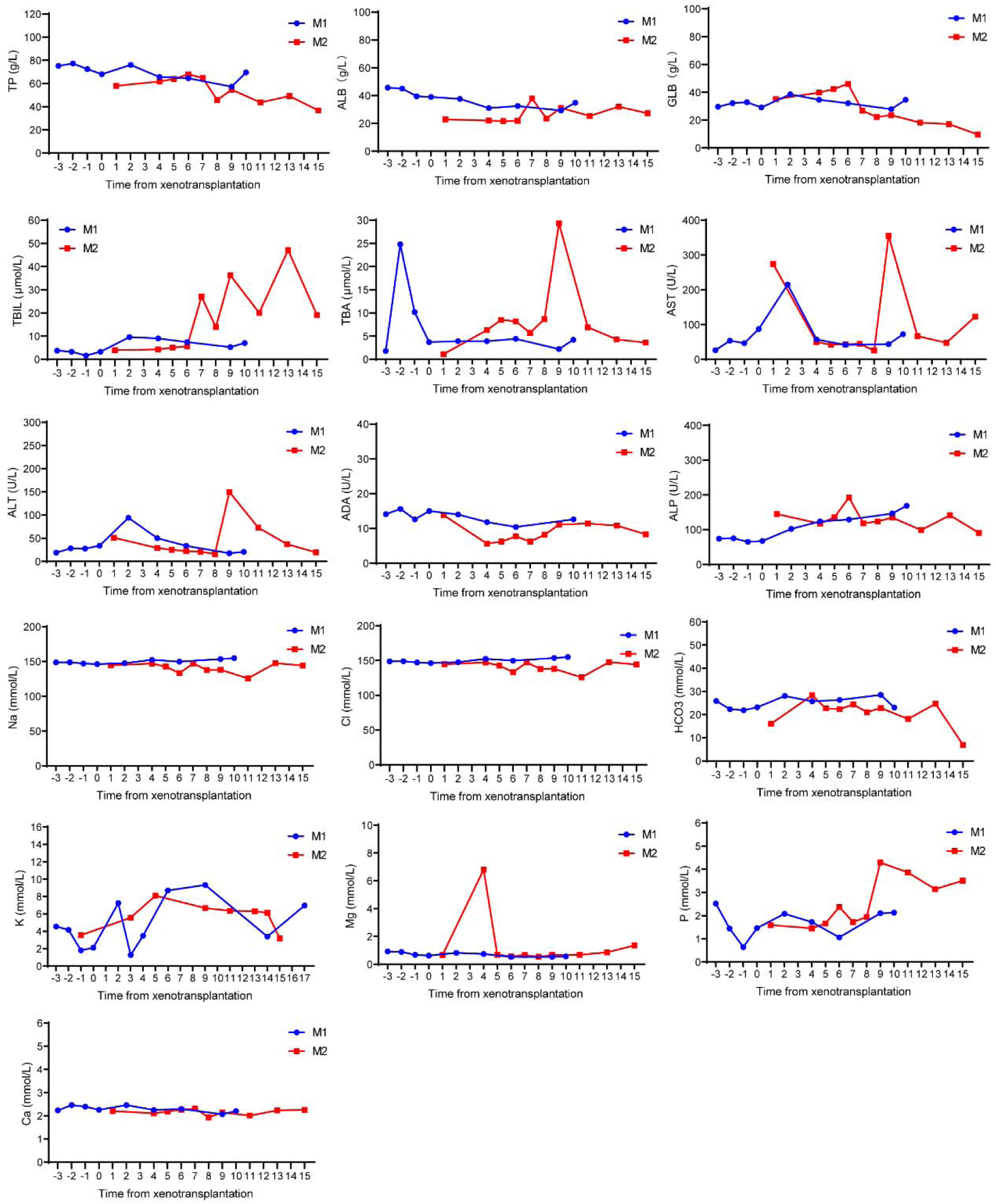
The liver functions indexes and electrolytes of recipient monkey. Liver function indexes: total protein (TP), ailbumin (ALB), and gloubulin (GLB), total bilirubin (TBIL), total bile acid (TBA, aspartate transaminase (AST), alanine transaminase (ALT), adenosine deaminase (ADA) and alkaline phosphatase (ALP); Electrolytes: sodium (Na), chlorine (Cl), bicarbonate (HCO3), calcium (Ca), magnesium (Mg), phosphorus (P) and potassium (K).

## DISCUSSION

Pig-to-human xenotransplantation is about to enter clinical phase, thus it is essential to construct and select donor pigs with most suitable genetic combination. The numbers of multigene-modified donor pigs for xenotransplantation are increasing with the advent of gene editing technologies. However, which gene combination is suitable for which organ or tissue transplantation remained unclear, and as larger the number of gene edited, as much difficult, and more time consuming to obtain viable donor pigs. Keeping in view, this study used the CRISPR/Cas9 system, PiggyBac transposase system and somatic cell cloning technology to obtain 8-GEC pigs and verified its effectiveness by transplanting the 8-GE kidney to NHP, thereby providing clinical evidence for pig-to-human kidney transplantation, and laid down the foundation for xenotransplantation research.

In order to produce 8-GEC pigs, we integrated 7 sgRNAs into one plasmid to achieve simultaneous KO of multiple porcine endogenous genes. This method is more effective than co-transfection of 7 plasmids where simultaneous transfer of multiple plasmids is difficult, and also there are more chances of cytotoxicity to cells. Recent research has also shown that this method can achieve efficient simultaneous editing of multiple genes (Zhang et al., 2022). Among eight randomly selected cell colonies, the GGTA1 gene editing efficiency was approximately 100%. On the premise that the WT genotype did not appear in the GGTA1 gene, the colonies for other two genes were further identified, and finally a cell line with a simultaneous editing of total of 13 alleles of 8 genes was selected. However, there were three genotypes of the GGTA1 gene in this cell line, which may be due to impure cell colonies. Therefore, we obtained the fetus through the first round of somatic cell cloning and conducted genotyping, and gene expression identification and karyotyping analysis to determine that it was a normal fetus with 8-GE. We then recloned to obtain 8-GEC pig and the proportion of healthy piglets among the cloned pigs was as high as 82% (23/28), after recloning. This once again proved that the method of first cloning to obtain fetuses, identifying the fetuses, and then re-cloning to obtain the piglets has effectively improved the production efficiency of GEC pigs (Zhao et al., 2020). In addition, the integrated sequence length for plasmids of hCD46, hCD55, hCD59, hTBM, hCD39, promoters, reached up to 14kb, and the piggyBac transposons system was used for transgenes to achieve efficient large fragment integration (64%, 16/25), which is similar to our previous study (Yue et al., 2021). Moreover, the copy number of PiggyBac-mediated gene integration remained stable in various organs and tissues of cloned piglets, suggesting that the cloning process or embryonic development did not affect the gene copy number.

In the 8-GEC pigs, the expression of hCD46, hCD55, hCD59, hTBM and hCD39 in different tissues was detected. However, the expression levels of each genes were not consistent among different individuals cloned from the same fetal cell line, which could be due to reprogramming during cloning. Moreover, we found that genes with high mRNA expression levels were not equally expressed at the protein level. For example, hCD59 which has low mRNA levels in porcine kidneys, while protein levels were higher (Figure 4). Moreover, the expression of the same gene (hCD59) is also different in different tissues of the same cloned pig (Figure S1), which may be due to different degrees of epigenetic modification of the human genes in the pig genome (Alhaji et al., 2019). Despite this, the pig kidneys removed after loss of function still exhibited high expressions all human genes hCD46, hCD55, hCD59, hTBM, and hCD39 as shown by immunofluorescence staining (Figure S3E).

In pig-to-monkey kidney transplantation, the 8-GE pig kidney showed good anti-immune rejection effects. First, *in-vitro* cross-matching experiments revealed that 8-GEC PBMCs had significantly reduced monkey IgG and IgM binding abilities, and increased anti-complement-dependent cytotoxicity. Secondly, during the 9 days post-operative observation, the immunoglobulins IgG, IgM and IgA as well as complements C3 and C4 remained relatively stable in the recipient monkeys, suggesting that the transfer of 3KO and three complement regulatory proteins effectively protected against the hyperlipidemia and acute rejection. However, at the experimental end point, the number of monocytes in the recipient monkeys increased significantly. Immunohistochemical analysis revealed that obvious infiltrates of CD57+ NK cells and CD68+ macrophages in the pig kidney grafts, suggesting that acute rejection occurred later. However, it has been reported that acute rejection is reversible, and timely adjustment of the dosage of immunosuppressants can help recipient monkeys to overcome the risk period of acute rejection (Cooper, 2020). Additionally, on the basis of 8-GE pigs, further transfer of hCD47 can also effectively inhibit acute rejection mediated by NK cells and macrophages (Nomura et al., 2020). In terms of anticoagulation, PT, TT and FIB remained relatively stable, and only APTT showed an increasing trend in the late transplantation period, indicating that hTBM and hCD39 overexpression effectively controlled coagulation. However, near the end of the experiment, the number of platelets dropped sharply, which may be related to the use of anti-inflammatory drugs (Bikhet et al., 2022).

Within 10 days after surgery, the 8-GE pig kidney functioned normally. The recipient monkey’s serum creatinine, cystatin C, uric acid, and urea nitrogen were maintained at relatively stable levels, and the 24-hr urine output was above 400 mL. In addition, immunosuppressants, antibiotics and other drugs did not cause any obvious liver damage or electrolyte imbalance. Unfortunately, an autopsy revealed that the recipient monkey had gastric perforation, and food and medicines sieved through the abdominal cavity. The immunosuppressant blood concentration of the recipient monkey did not meet the standard levels (Figure S3-C), and also severe nutritional deficiencies might affect the survival time of the recipient monkey. Thus, we believe that improvements in postoperative care plans and experiences will be helpful in increasing the survival time of 8-GEC pig kidneys into non-human primates.

In summary, this study used the CRISPR/Cas9 system, PiggyBac transposase system and somatic cell cloning technology to successfully obtain 8-GEC donor pigs, and proved that 8 gene editing effectively alleviates the immune incompatibility in xenografts as evidenced by survival of the kidney xenograft from these pigs to NHP.

## MATERIALS AND METHODS

Experimental animals used in this study and all surgical procedures were approved by the Animal Ethics Committee of Yunnan Agricultural University.

### Vector construction

To target porcine GGTA1, CMAH and β4GalNT2, 2∼3 sgRNAs were designed at the 3^rd^ exon, 4^th^ exon and 2^nd^ exon of these three genes, respectively. All the sgRNAs were expressed using the U6- promoter and subcloned into the PX458 vector (Addgene ID: 112220) to construct the PX458-sgRNAs targeting vector. Meanwhile, PB-hICAM2-hTBM-P2A-hCD39-hEF-1α-hCD46-P2A-hCD55-P2A-hCD59-P2A-Pur o vector driven by human ICAM2 promoter (hTBM and hCD39), and the human EF-1α promoter (hCD46, hCD55 and hCD59) was constructed and was confirmed by sequencing, followed by plasmid extraction and cell transfection.

### Cell transfection, screening, and identification

The *Diannan* miniature pig fetal fibroblasts (PFFs) with O blood type were cultured into DMEM containing 10% fetal bovine serum (FBS) at 38°C and 5% CO_2_. Once the cell attained 60%confluence, the cells were harvested and counted. Approximately, 3×10^5^ cells were suspended in the electro-transfection buffer containing PX458-sgRNA plasmid (8μg), PB-hICAM2-hTBM-P2A-hCD39-hEF-1α-hCD46-P2A-hCD55-P2A-hCD59-P2A-Pur o plasmid (8μg), and transposase plasmid (4μg) were incubated for 5 min, and electroporated under the program of EH-100 by a Lonza 4D-nucleofector X Unit (EH-100, Germany). After electroporation, cells were plated into T25 flasks for 24 h in DMEM supplemented with 10% FBS. Then, 3μg/ml puromycin was added to the culture medium for 24−48 h to select successfully transfected cells. Subsequently, the survived cells were digested and ∼80 cells were seeded in a 100 mm diameter dish and cultured for 8 days to obtained single-cell colonies, which were further harvested for genotyping by PCR and Sanger sequencing using the primers shown in Table S1. Cells with biallelic knockout (KO) of GGTA1, CMAH, and β4GalNT2, and hICAM2-hTBM and -P2A-hCD39-hEF-1α-hCD46-P2A-hCD55-P2A-hCD59-Puro integrated colonies were selected as a donor cell for SCNT.

### Somatic cell nuclear transfer and embryo transfer

Oocyte collection, in vitro maturation, SCNT, and embryo transfer were performed as described previously (Wei et al., 2013). Briefly, cultured cumulus-oocyte complexes (COCs) were isolated from cumulus cells by treating with 0.1% (w/v) hyaluronidase.

The oocytes with first polar body were selected and cultured in PZM-3 media containing 10% FBS, 0.0171g/mL sucrose, and 0.01μg/mL colchicine and incubated at 38°C, 5 % CO_2_ incubator for 0.5∼1 h. The first polar body was enucleated via gentle aspiration using a beveled pipette in TLH-PVA, while the donor cells were injected into the perivitelline space of the enucleated oocytes. The reconstructed embryos were fused with a single direct current pulse of 25V/mm for 20 µs using the Electro Cell Fusion Generator (LF201, NEPA GENE Co., Ltd., Japan) in fusion medium. Embryos were then cultured in PZM-3 for 0.5–1 h and activated with a single pulse of 150 V/mm for 100 µs in activation medium. The embryos were equilibrated in PZM-3 supplemented with 5 µg/ml cytochalasin B for 2 h at 38.5°C in a humidified atmosphere with 5% CO_2_, 5% O_2_, and 90% N_2_ (APM-30D, ASTEC, Japan) and then cultured in PZM-3 medium with the same culture conditions described above until embryo transfer. The SCNT embryos were surgically transferred into the oviducts of the recipients, and a viable fetus was obtained through caesarian section after 33 days of pregnancy. A fetal fibroblast cell line was established, and subjected to PCR, T7EI enzyme digestion experiment, Sanger sequencing, flow cytometry, immunofluorescence, and WB identification. Fetuses with biallelic KO of 3 genes, and targeted overexpression of 5 genes were selected as donor cells for recloning. After completion of gestation period (114 days), GE cloned piglets were obtained through natural delivery and identified by the same way of fetus identification.

### Genotype identification of cell colonies, cloned fetuses and piglets

The screened cell colonies were directly lysed, and the whole-genome DNA from cell colonies, fetal tissue and cloned piglet ear tissues was extracted using high-salt method and used as a template for the amplification of the target gene fragment through PCR reaction. For cell colonies, we first confirmed the integration of five genes through gel electrophoresis imaging, and then used Sanger sequencing analysis to determine the genotypes of the target regions of three gene GGTA1, CMAH, and β4GalNT2, and finally the cells with simultaneous knock-out (KO) of three genes and successful integration of five genes were selected as a donor for cloning. The 8-GE status of fetuses and cloned piglets were analyzed by PCR and Sanger sequencing.

### Karyotyping

PFFs at 70%∼80% confluency were incubated with 0.2 μg/mL colchicine (Colcemide-Gibco etc.) for 2 h, and treated with hypotonic solution Na-Citrate/NaCl at 37°C for 15 min. Subsequently, cells were fixed with methanol/acetic acid glacial (3:1) for 20 min at -20°C and washed twice with methanol/acetic as stated above, and Gimsa stained according to manufacturer guidelines and analyzed for metaphase chromosome smear analysis using SmartType.

### Droplet digital PCR (ddPCR) for transgenic copy numbers

The samples including the human blood (positive control), the pre-transfected porcine cells (negative control), the selected cell lines, the fetal cells, and the heart, liver, lung, kidney, spleen and other tissues of the cloned pig derived from fetuses were collected and extracted DNA. Then, these DNA samples were digested by MseI (Cat#FD0984, ThermoFisher), EagI (Cat#FD0334, ThermoFisher) and AsisI (Cat#FD2094, ThermoFisher) restriction endonuclease at 37°C for 2 h, and inactivated at 65°C and 85°C for 20 min. The digested product was diluted to 5ng/ul and used as a ddPCR template. The primers and probes corresponding to hCD46, hCD55, hCD59, hTBM, hCD39 and GAPDH genes (Table S1), ddPCR super mix (Cat#1863024, Bio-Rad), and DNA template were mixed to prepare a 25uL reaction system. Then, ddPCR droplets were generated with QX100 Droplet Generator (Bio-Rad) and transferred to a 96 well plate. PCR reactions were performed at 94°C for 5 min, 94°C for 30 s, 56°C for 1 min (40 cycle), and 98°C for 10 min. After the reaction, the 96 well plate was placed in Bio-Rad Droplet Reader and QunataSoft Software was used to set up the experimental design and read the experiment. Once the program was finished, the copy numbers were analyzed through gating, according to the manufacturer’s instructions (Bio-Rad).

### Quantitative polymerase chain reaction

In order to evaluate the mRNA expression levels of hCD46, hCD55, hCD59, hTBM, and hCD39, total RNA from heart, liver, lung, kidney and other tissues of WT and GEC pigs, as well as human umbilical cord tissues, was extracted using TRIzol reagent (Cat#ET111-01-V2, TransGen Biotech) according to the manufacturer’s instructions, and the concentration and RNA quality were detected. Complementary DNA (cDNA) was synthesized from total RNA using a PrimeScript RT reagent Kit (Cat#RR047B, TaKaRa) and was used as a template to perform qPCR in TB green-based qPCR instrument (CFX-96, Bio-Rad, USA). The reaction was performed in a 20 µl reaction mixtures comprising 10 µl of 2×TB Green® Premix Ex Taq™ (Tli RNaseH Plus) (Cat#RR420A, TaKaRa), 1 µl of cDNA, 1 µl of forward primer, 1 µl of reverse primer, and 7 µl of ddH_2_O (primers listed in Table S1). The reaction program is as follows: 95°C for 30 s, followed by 40 cycles of 95°C for 10 s, and 62°C for 45 s. Three technical replicates were conducted for each sample and the relative expression levels of target genes were quantified by 2^-ΔΔct^.

### Immunofluorescence

The paraffin-embedded tissue blocks were cut into 5 µm, transferred to glass slides, dewaxed using xylene and gradient alcohol, put into a microwave oven to retrieve antigens with EDTA buffer (Cat#G1207, Servicebio Bio) at 92–98°C for 15 min, and cooled at room temperature. Then, sections were washed with PBS for three times (each time 3 min), incubated with autofluorescence quencher A (Cat#G1221, Servicebio Bio) at RT in dark for 15 min, washed with PBS for three times again, incubated with FBS at RT for 30 min, and dried. The dried sections were incubated with corresponding antibodies (Table S2). For visualization, corresponding secondary antibodies (Table S2) were diluted with PBS containing 10% FBS (v/v = 1:200) and used to incubate sections at 4°C in dark for 2 h, and a negative control was incubated with PBS containing 10% FBS. Then, sections were washed with PBS three times and stained with DAPI (Cat#G1012, Servicebio Bio) for 3 min. After washing with PBS for 1 min, autofluorescence quencher B (Cat#G1221, Servicebio Bio) was added for 5 min and washed three times again. Finally, sections were mounted with anti-fluorescence quencher (Cat#G1401, Servicebio Bio) and imaged using an OLYMPUS BX53 fluorescence microscope, and the fluorescence intensity of different samples was compared using ImageJ software.

### Pig-monkey cross-matching experiment

Pig PBMCs and rhesus monkey serum were used for cross-matching to select recipient monkeys with low antibody titers and low complement-dependent toxicity for kidney transplantation.

Whole blood (10 mL each) of healthy rhesus monkeys was collected and serum was separated. At the same time, 10 mL of whole blood from wild-type pigs and gene-edited donor pigs were collected to isolate PBMCs. Based on the antigen-antibody binding assay, inactivated serum from each monkey was incubated with wild-type pig and gene-edited donor pig PBMC, and then treated with anti-IgG antibody (Cat#628411, Invitrogen) and anti-IgM antibody (Cat#A18842, Invitrogen), respectively), and the levels of binding of IgG and IgM to pig cells was detected by cell flow cytometry, and monkeys with lower MFI were selected as xenograft recipients.

For complement-dependent cytotoxicity experiment, 50 uL of 50% inactivated monkey serum was incubated with pig PBMC cells for 2 hr. The reaction was then terminated using 0.5 mL pre-cooled PBS, samples were centrifuged at 300×g for 3 min, the supernatant was discarded, and incubated with 50 uL rabbit complement serum (1: 10, Cat#S7764, Sigma) at room temperature for 30 min. Finally, cells were stained with PI staining solution (Cat#GA1174, Servicebio Bio) for 10 min, and analyzed for cell death using a CytoFLEX flow cytometer (Beckman Coulter, USA).

### Pig-to-monkey kidney xenotransplantation and postoperative care

We carried out pig-to-nonhuman primate kidney transplantation at Yunnan Province Key Laboratory for Porcine Gene Editing and Xenotransplantation, Yunnan Agricultural University, Kunming. On the day of surgery, both kidneys of the recipient monkey were removed, an 8-GE pig kidney was transplanted, and a gastrostomy was performed to facilitate postoperative medication. After the operation, urine biochemistry, blood biochemistry, immune indicators, coagulation function, infection, immunosuppressant blood drug concentration, etc. were tested regularly. The blood flow of the transplanted kidney was detected by ultrasound. Clinical, and medical staff and veterinary professionals provided postoperative cares, including various tests, indicators and the mental state of the recipient monkey, and apply corresponding treatment methods.

### H&E staining

Tissue samples were fixed in 4% paraformaldehyde for 48–72 h, processed by an automatic tissue processor (Yd-12p, Jinhua Yidi, China) and embedded in a paraffin block (Yd-6D, Jinhua Yidi, China). The paraffin blocks were cut into 3-μm-thick sections using a Microm HM 325 microtome (Thermo Scientific, USA) and allowed to dry on glass slides overnight at 37°C. Thereafter, the tissue sections were deparaffinized in xylene and rehydrated through graded ethanol dilutions. Sections were stained with hematoxylin–eosin (H&E) (Cat#G1120, Solarbio) according to manufacturer’s instruction. Imaged using OLYMPUS BX53 fluorescence microscope and analyzed using software of accessories.

### Immunohistochemical analysis of xenografts

The paraffin-embedded tissue blocks were cut into 3 µm, transferred to glass slides, dewaxed using xylene and gradient alcohol, put into a microwave oven to retrieve antigens with EDTA buffer (Cat#G1207, Servicebio Bio) at 92–98°C for 15 min, and cooled at room temperature. Then, sections were washed with PBS for three times (each time 3 min), incubated in 3% H_2_O_2_ solution for 15 min, washed with PBS for three times again, incubated with FBS at RT for 30 min, and dried. The dried sections were incubated with corresponding antibodies (CD57, 1:200; CD68, 1:200, Table S2; Table S2) at 4°C overnight After washing with PBS for thrice, sections were incubated with 5 μg/mL HRP-conjugated goat anti rabbit/mouse IgG antibody for 20 min. After washing thrice again, sections were stained with fresh DAB (KIT-9901, Elivision TM plus Polyer HRP IHC Kit, China) solution in dark for 1 min. PBS wash 3 min × 3. Hematoxylin counter-staining, and neutral gum sealing slides. Imaged using OLYMPUS BX53 fluorescence microscope and analyzed using software of accessories.

### Statistical analysis

All data were analyzed using SPSS 22.0 software package (IBM Crop, Armonk, NY) and expressed as mean ± standard deviation of mean (SD). Statistical significance was defined as *P < 0.05, **P < 0.001.

## DATA AVAILABILITY STATEMENT

The data generated in the present study may be shared on reasonable request from the corresponding author.

## Compliance and ethics

The Author(s) declare that they have no conflict of interest.

## Acknowledgments

We thank the Ministry of Science and Technology of the People’s Republic of China (Grant No. 2019YFA0110700) and the Department of Science and Technology of Yunnan Province (Grant No. 202102AA310047) for their financial support. We extend our thanks to Professor Chen Gang (Tongji Hospital Affiliated to Tongji Medical College of Huazhong University of Science and Technology) for providing surgical intervention, postoperative care and guidance; and to Lai Ren (Researcher, Kunming Institute of Zoology, Chinese Academy of Sciences) for providing CVF.

## Authors’ Contributions

HJW conceived and designed the experiment, HYZ and ZZ designed the research. JW, KX, HZ, MAJ, GX, XH, and CY performed the molecular experiments and analyzed data. DJ, HZ transfected the cell and performed a cell culture. JG, TW, KL, and XZ carried out SCNT and embryo transfer. FW, GW, and HZ performed pig to monkey kidney xenotransplantation. KX, MAJ wrote the manuscript. HJW, HYZ, ZZ, and MAJ revised the manuscript and validated the data. All authors contributed to the article and approved the submitted version.

---

**Figure S1:**
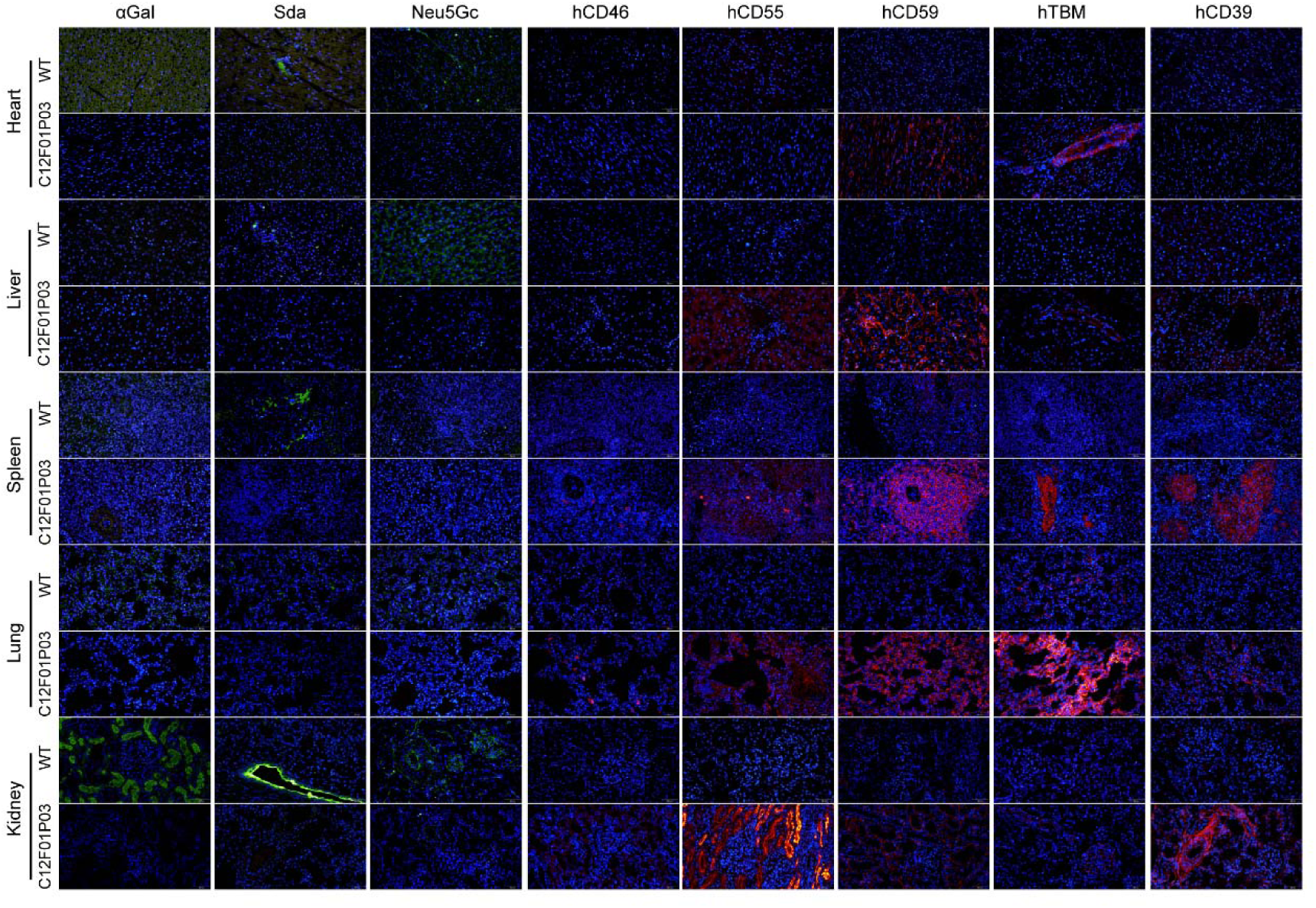
Immunoflourescence staining of different tissues of 8-GEC piglets. The protein expression of αGal, Neu5Gc, Sda, hTBM, hCD39, hCD46, hCD55 and hCD59 genes in kidney, lung, liver, spleen and heart of cloned piglets was confirmed by immunofluroscence.

**Figure S2:**
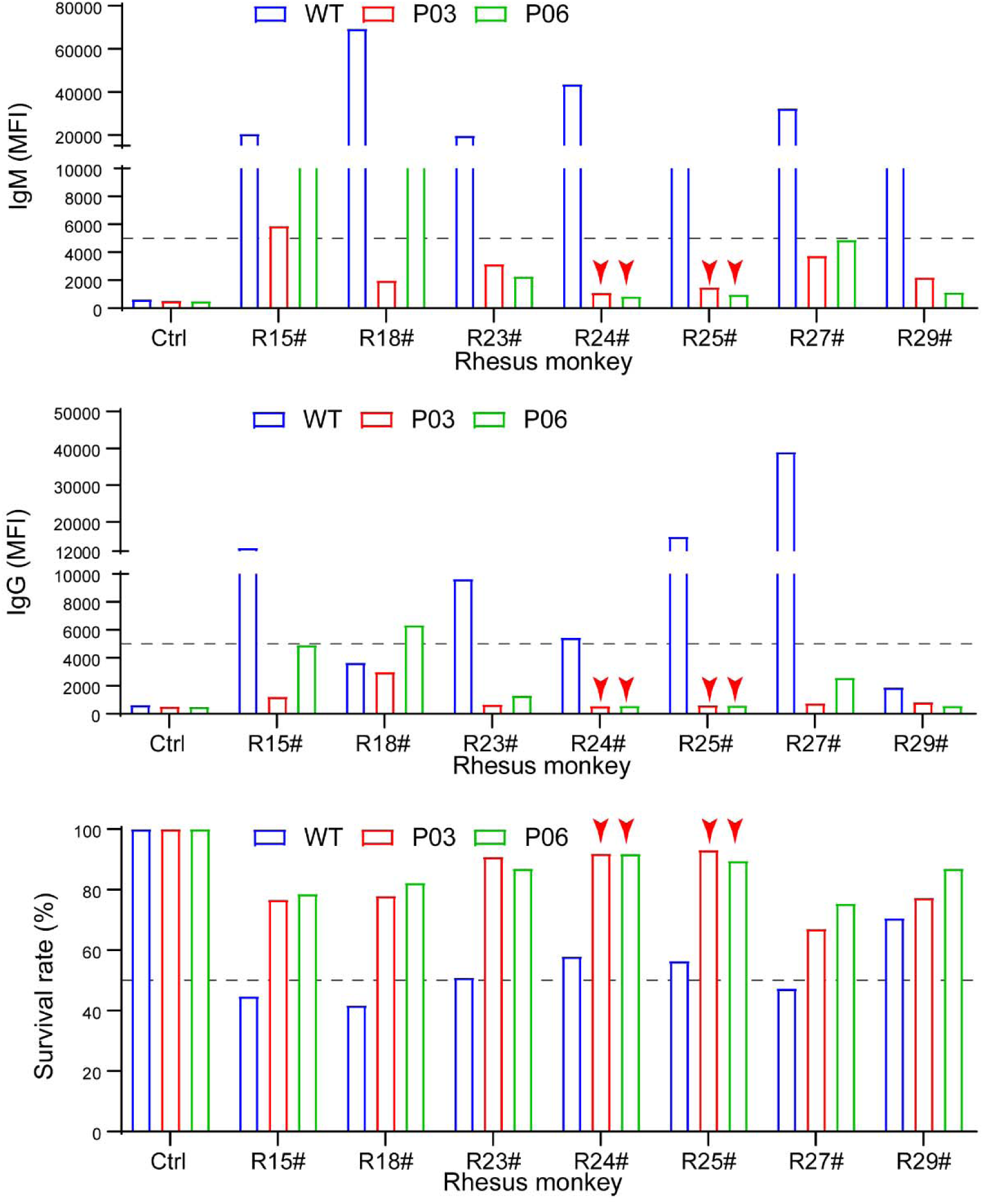
Crossmatch between the 8-GEC pigs and rhesus monkeys. **A**. The levels of monkey IgM binding to 8-GEC porcine peripheral blood mononuclear cells (PBMCs). **B**. The levels of monkey IgM binding to 8-GEC porcine PBMCs. **C**. The survival rate of 8-GEC porcine PBMCs after 50% monkey serum incubation for 2 hours.

**Figure S3:**
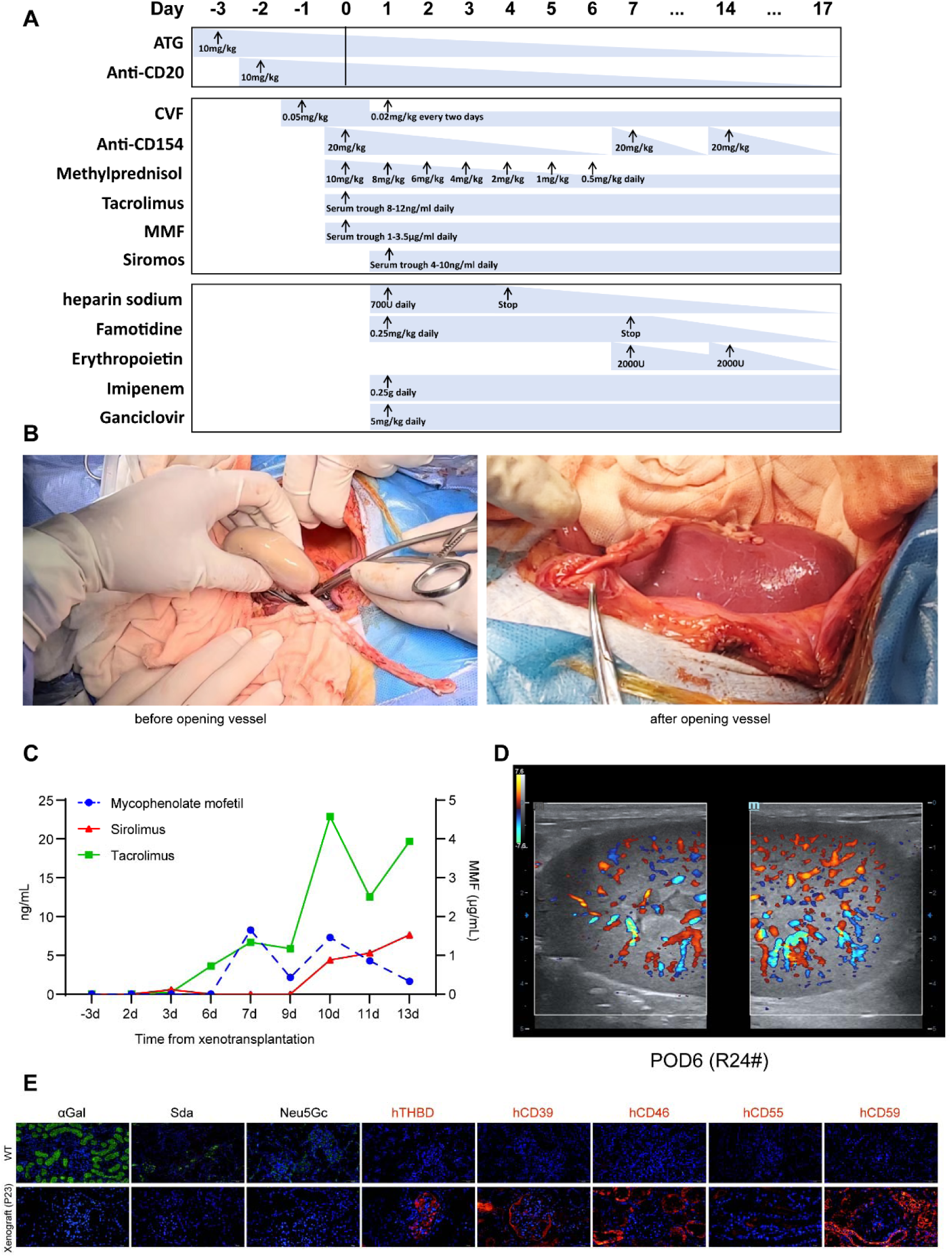
Immunosuppressive regimen and other observations. **A.** Immunosuppressive regimen, **B.** Changes of pig kidney before and after opening the blood vessel during transplantation into rhesus monkeys **C.** Measurement of immunosuppressant concentrations in the blood. **D.** Doppler ultrasonography of pig kidney xenograft after 6 days of surgery. **E.** Expressions human genes after kidney function loss as showen by immunofluorescence staining

**Table S1.**
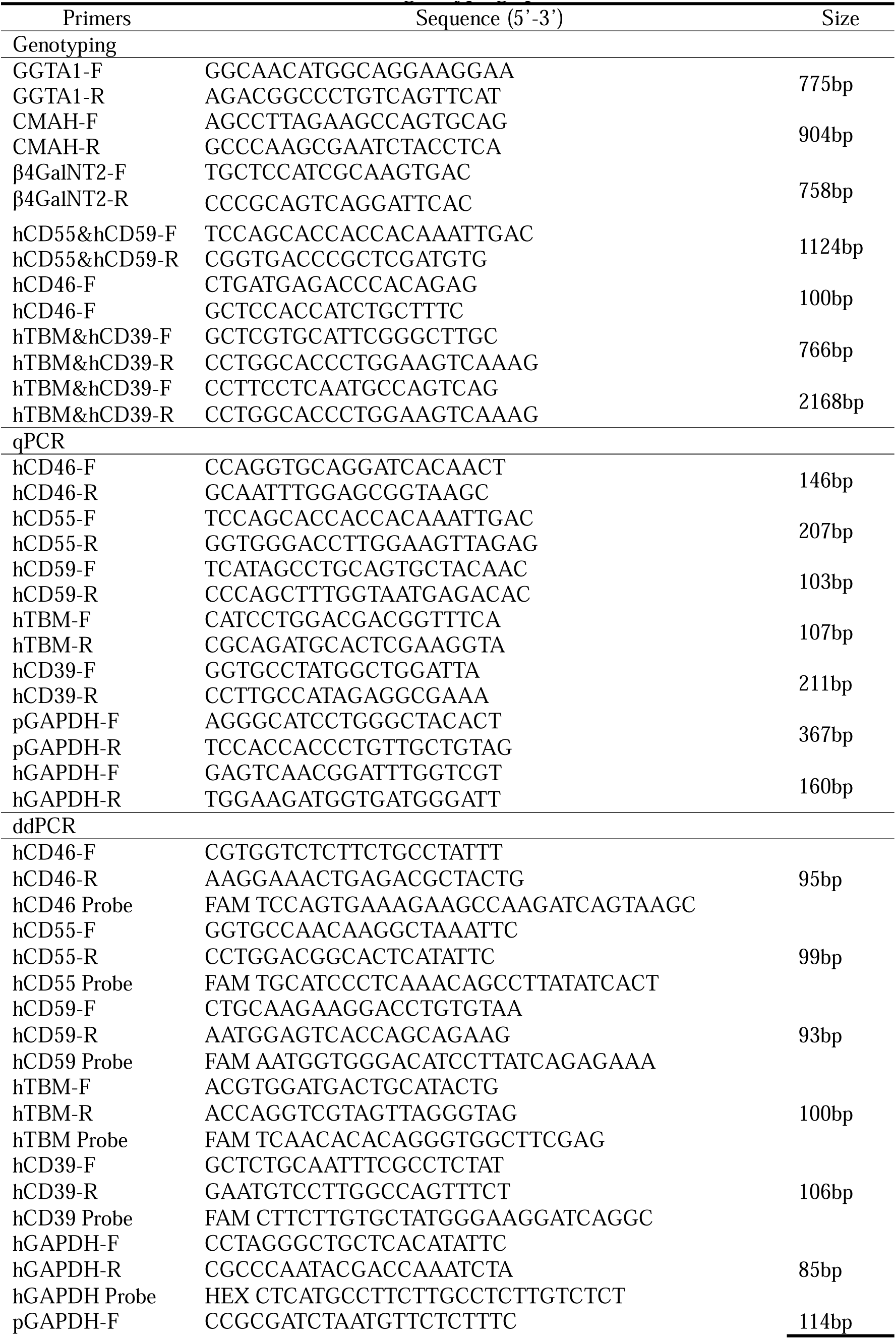
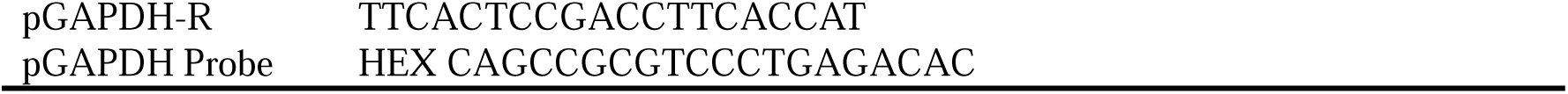
Primers for genotyping, qPCR and ddPCR.

**Table S2.**
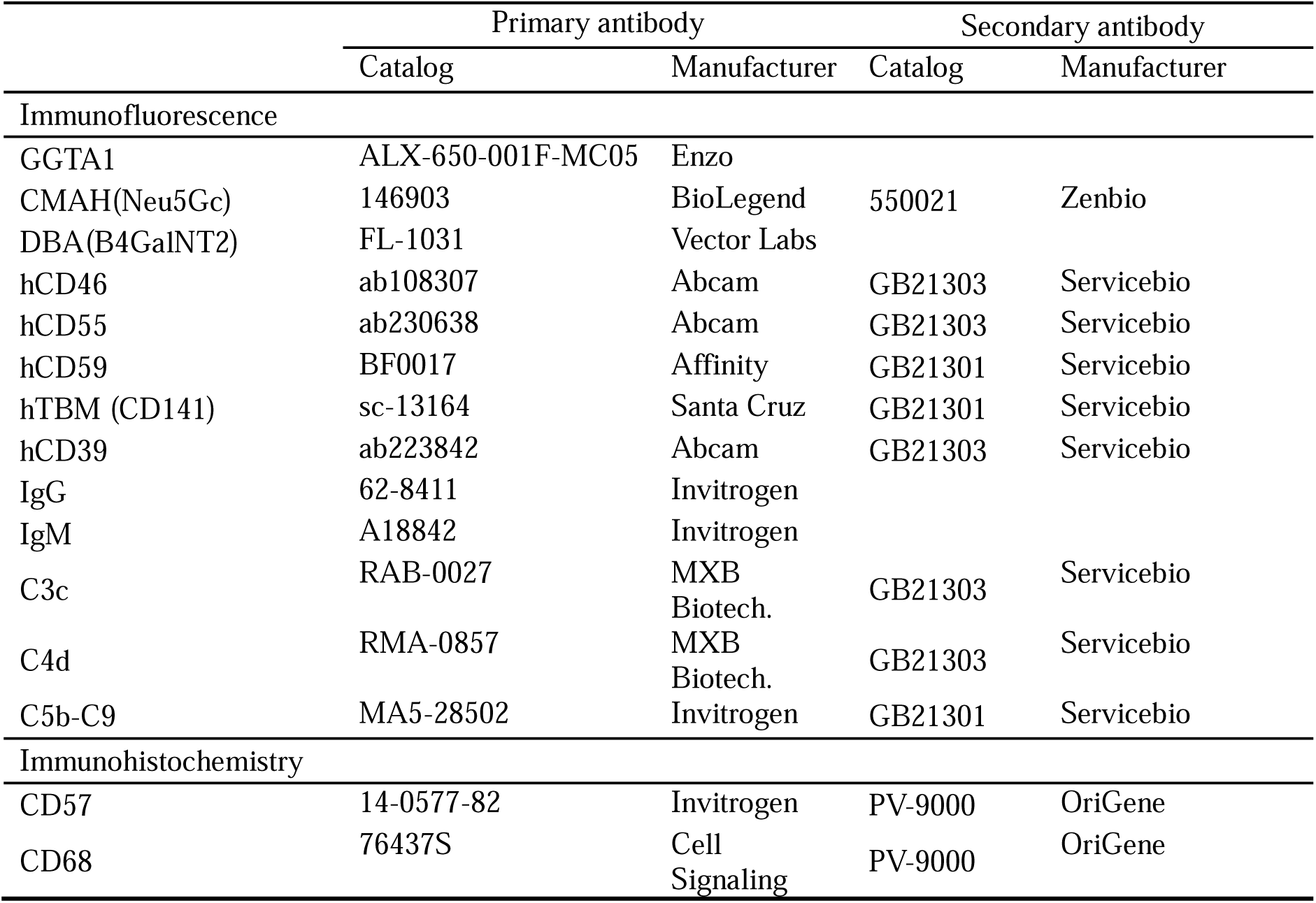
A summary of antibodies used by immunofluorescence and immunohistochemistry.

